# Rescuing Off-Equilibrium Simulation Data through Dynamic Experimental Data with dynAMMo

**DOI:** 10.1101/2023.05.23.541878

**Authors:** Christopher Kolloff, Simon Olsson

**Affiliations:** Chalmers University of Technology, Department of Computer Science and Engineering, Rännvägen 6, 412 58 Gothenburg, Sweden

## Abstract

Long-timescale behavior of proteins is fundamental to many biological processes. Molecular Dynamics (MD) simulations and biophysical experiments are often used to study protein dynamics. However, high computational demands of MD limit what timescales are feasible to study, often missing rare events, which are critical to explain experiments. On the other hand, experiments are limited by low resolution. We present dynamic Augmented Markov models (dynAMMo) to bridge the gap between these data and overcome their respective limitations. For the first time, dynAMMo enables the construction of mechanistic models of slow exchange processes that have been not observed in MD data by integrating dynamic experimental observables. As a consequence, dynAMMo allows us to bypass costly and extensive simulations, yet providing mechanistic insights of the system. Validated with controlled model systems and a well-studied protein, dynAMMo offers a new approach to quantitatively model protein dynamics on long timescales in an unprecedented manner.

## Introduction

Understanding the triad of protein structure–function–dynamics is of paramount importance in many fields, including biochemistry, biophysics, and medicine [1–7]. Thanks to extensive studies of the bovine pancreatic trypsin inhibitor (BPTI), for example, we now understand the atomic details of its essential role in inhibiting serine proteases [8]. This knowledge has been possible by combining findings from the fields of X-ray crystallography [9] and Nuclear Magnetic Resonance (NMR) [10–13] with Molecular Dynamics (MD) simulations [14–16]. However, reconciling experimental and simulation data in a systematic manner often poses problems due to technical and resource limitations. Enabling such a merger would yield a significant opportunity for quantitative structural biology and biophysics.

Typically, we model dynamic experiments, [17] such as NMR relaxation dispersion, single molecular Förster Resonance Energy Transfer (FRET), dynamic neutron scattering, or X-ray Photon Correlation Spectroscopy using simple *n*-site jump models [18–22]. This approach yields forward and reverse exchange rates for the different states as well as site populations. However, modeling dynamics this way beyond a two-state exchange is challenging, due to experimental limitations and poor timescale separation. In effect, we are limited to highly simplified models of the complex underlying dynamics of our data where the structure of states often remain elusive or ambiguous [23–28].

Over the past few decades, molecular dynamics (MD) simulations have become increasingly popular in the field of biophysics, providing atomistic insights into the behavior of biological systems at high temporal and spatial resolutions [29–31]. Force field models are steadily improving in quality and their scope is boarding to include disordered proteins and nucleic acids [32–34]. Although not broadly available, the development of special-purpose computers, like Anton [35–37], makes it possible to study millisecond timescale molecular processes. GPU-accelerated simulations as well as distributed computing initiatives, such as Folding@home [38] or GPUGrid are more widely available allow us to access processes on the micro-to millisecond timescale, in particular when large ensembles of simulations are analyzed using Markov state models (MSM) [39–43]. MSMs represent the *molecular kinetics* fully: the relevant structural states, their thermodynamic weights and their mutual exchange rates [44–50]. MSMs have enabled studies of many biological processes, such as protein folding, enzymatic activity, or protein–protein interactions [42, 51–57].

Despite these advances in force field accuracy and simulation technologies, we often observe a systematic discrepancy between the experimental values predicted from Molecular Dynamics (MD) trajectories and experimental data [58–61]. The origin of these discrepancies is two fold. First, imperfections in the force field models remain which lead to skewed populations and altered dynamics. Second, simulations still do not cover the range of biological timescales of interest. Methods like transition [62] and discrete path sampling [63], transition interface sampling [64], milestoning [49, 65], metadynamics [66], flooding [67], or replica exchange [68, 69] offer potential avenues to bridge this timescale gap. The advent of deep generative neural network-based surrogates further provides new venues for overcoming the sampling problems [70, 71]. However, each of these approaches have inherent limitations in their scope or rely on extensive manual intervention. Together, these limitations prevent us from directly comparing to many experiments and thus gaining a mechanistic interpretation of our data.

The integration of simulation and experimental data is a big challenge with a long history [72]. Previous methods include *post hoc* reweighing or sub-selection of simulations data [73, 74], modeling kinetics with generated ensembles [75], biasing simulations with stationary experimental data [76–85] and dynamic experimental data [86, 87], and building Markov state models using experimental and simulation data [88], Augmented Markov models. Augmented Markov Models (AMMs) combine stationary experimental observables with simulation data to correct for bias in MD data, which improved agreement with complementary stationary and dynamic data [88]. However, AMMs cannot take into account dynamic data, and consequently also cannot deal with situations where our simulations have not sampled processes which are important to explain the data.

Here, we present dynamic Augmented Markov Models (dynAMMo), a new approach that accounts for stationary and dynamic experimental data, such as *R*_1*ρ*_ or Carr-Purcell-Meiboom-Gill (CPMG) relaxation dispersion experiments [20, 89, 90], to estimate a Markov model (Figure 1). By combining constrained optimization with the principle of maximum entropy, we are able to correctly recover experimental timescales from biased simulations and are able to model exchange between states not seen in the simulation data as long as the states themselves are known. To our knowledge, this is the first method that enables the construction of mechanistic models of protein dynamics, even when rare events remain unobserved in the MD data. This achievement is made possible through the dynamic experimental data that complement the simulations by reporting on the exchange processes that have not been sampled by the simulations. With dynAMMo, we therefore address the aforementioned issue in MD simulations by circumventing the, often very costly, need for (reversibly) sampling rare events in order to establish a kinetic mechanistic model. Also, we can show that our method fails and thus does not overfit when one or more relevant states are missing. The method therefore broadens the scope for future research in understanding the complex dynamics of biomolecular systems in general and brings the field closer to the development of data-driven models that accurately capture the underlying mechanisms-of-action.

**Figure 1:**
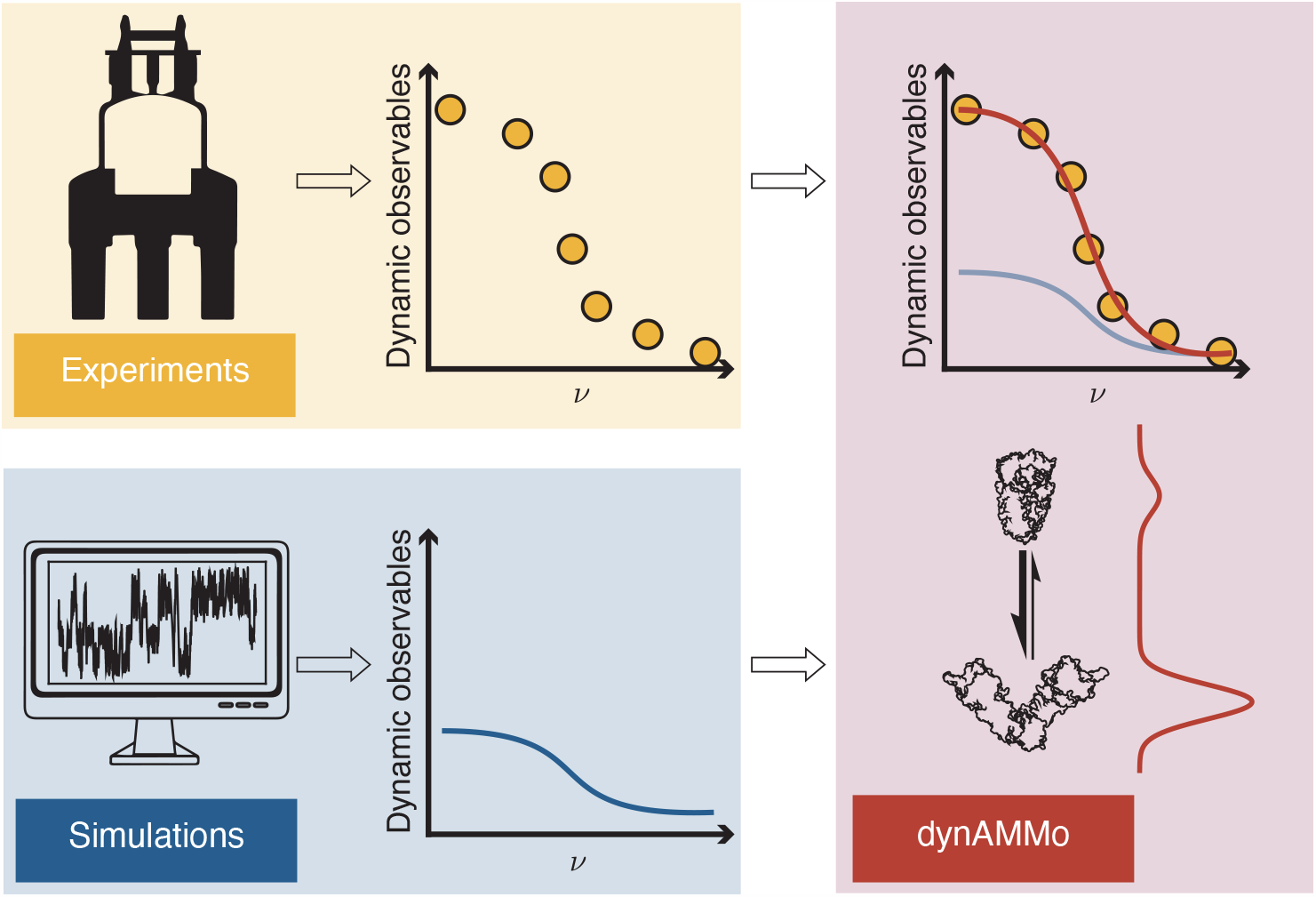
Schematic of dynamic Augmented Markov Models (dynAMMo). Simulations often fail to reproduce experimentally observed dynamic observables, such as NMR relaxation dispersion data due to force field inaccuracies and finite simulation length. Experiments, on the other hand, are often noisy and often do not reveal the mechanistic details of kinetic exchange. With dynamic Augmented Markov Models, we take into account, both, simulation and experimental data to obtain a single mechanistic kinetic model.

## Theory

### Dynamic Augmented Markov Models

Markov state models are based on the discretization of the state space Ω of a molecule into *n* states. By following the traversal of a MD trajectory through these states we can estimate the transition probabilities *p*_*ij*_ from states *i* to states *j* through the analysis of transition counts *c*_*ij*_(*τ*) with a lag time *τ* [39, 40, 43]. The resulting transition matrix **T**(*τ*) ∈ R^*n*×*n*^ encodes the *molecular kinetics* of the system, including the populations of the states and the rates of exchange between them. This information is accessible through the spectral components of *T* (*τ*), the eigenvectors **R** and eigenvalues ***λ***, as well as the stationary distribution ***π*** (see Supporting Information, ‘Theory’) [43].

Augmented Markov Models [88] aim to estimate a MSM which matches stationary experimental observables, such as NMR ^3^*J* -coupling or Residual Dipolar Coupling (RDC) data that probe the “true” Boltzmann distribution, by reweighing the relative populations of the states through the maximum entropy principle. Even though this approach does not directly take into account information about the kinetic rates between the states, Olsson and Noé observe that the integration of stationary observables has an effect on the prediction of dynamic observables, such as 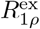 relaxation dispersion. However, in general we cannot expect AMMs to match dynamic experimental data, nor can they consider cases in which not all states of the MSM are connected.

Here, we address these limitations with dynamic Augmented Markov models that combine simulation data, in the form of one or more count matrices C (Supplementary Information, algorithm 1, line 1), and dynamic and stationary experimental observables **o**^exp^ to a single kinetic model. By combining these sources of information, we aim to obtain a more accurate and comprehensive representation of biomolecular dynamics. Using experimental data that report on the conformational exchange kinetics, we can directly estimate the forward and reverse rate constants of switching from one state to another. Unlike Brotzakis et al., no prior knowledge of the kinetic rates are required nor are we limited to a two-state exchange.

### Connection between experiments and simulations

For each experimental observable **o**^exp^, we assume that there is a corresponding observable function *f* (·) (*forward model*) available that maps all configurations, **x** ∈ Ω, to a complex or real vector space, V, however, often just a scalar, e.g., a distance or a chemical shift. For MSMs, we can average these values over the *n* Markov states yielding **a** ∈ *V*^*n*^ [88]. Here, we focus on dynamic experiments, where we measure time correlations of these observables either directly or through a transformation. From an MSM, **T**(*τ*), we can compute the time correlation of *f* :

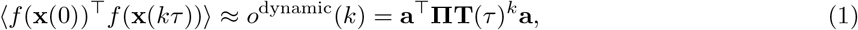

where **Π** is the diagonal matrix of the stationary distribution ***π***, and ^⊤^ is the transpose or the complex conjugate. We can compare this quantity directly to experimentally measured counterparts and thereby use it to drive the estimation of MSMs. Many dynamic observables, however, are transformations of the time-correlation, rather than the time-correlation itself. This includes, among others, Carl-Purcell-Meiboom-Gill (CPMG) and 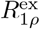 relaxation dispersion, which measure the convolution of the time-correlation function with a spin-lock field. Here we assume fast chemical exchange, and use previously described closed-form expression for Markov models [91, 92] to predict data and drive MSM estimation (see Supporting Information, ‘Theory’ for a more detailed explanation).

### Estimation of dynamic Augmented Markov Models

We estimate dynAMMo models by optimizing a loss function which includes the transition counts *c*_*ij*_ from states *i* to *j* and the sum of the mean-square difference between the predictions and experimental data, D, of the *l*th observable at the *k*th lag-time,

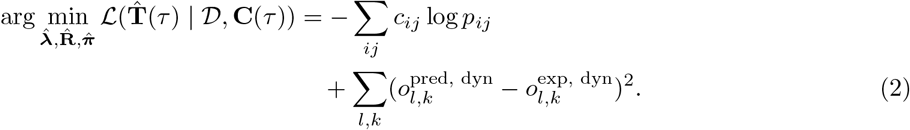

Here, *p*_*ij*_ is the probability of transitioning between states *i* and *j*. The loss is computed with respect to the spectrum of 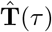, i.e., the eigenvalues 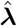, eigenvectors 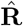, and the stationary distribution 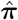 and is subject to several constraints. To estimate 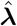 and 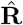, we use gradient descent with additional orthogonality constraints for optimization of the eigenvectors. Rather than enforcing orthogonality directly with a penalty term in the loss function, dynAMMo optimizes the eigenvectors on the Stiefel manifold through Riemannian optimization [93]. Following AMMs, we further have the option to include stationary experimental observables as described previously [88]. The estimation procedure as well as the theoretical details are explained in more detail in Supporting Information, ‘Theory.’

## Results and Discussion

### Enforcing dynamic experimental constraints on a kinetic model rescues biased simulation data

To demonstrate the power of dynamic Augmented Markov Models, we will first examine our model by applying it to two one-dimensional energy potentials: the Prinz potential [43] (Fig. 2, brown background) and the three-well potential (Fig. 2, teal background). In both model systems, there is one slow transition as well as one or more faster transition(s) that we aim to model. The Prinz potential has four states with comparable populations with one slow transition between the first and last two states, whereas the three-well potential has two fast-interchanging low-energy states and one state with a high energy barrier. In both model systems, we used the slowest eigenvectors as an observable function as they encode near perfect information about the slowest process.

**Figure 2:**
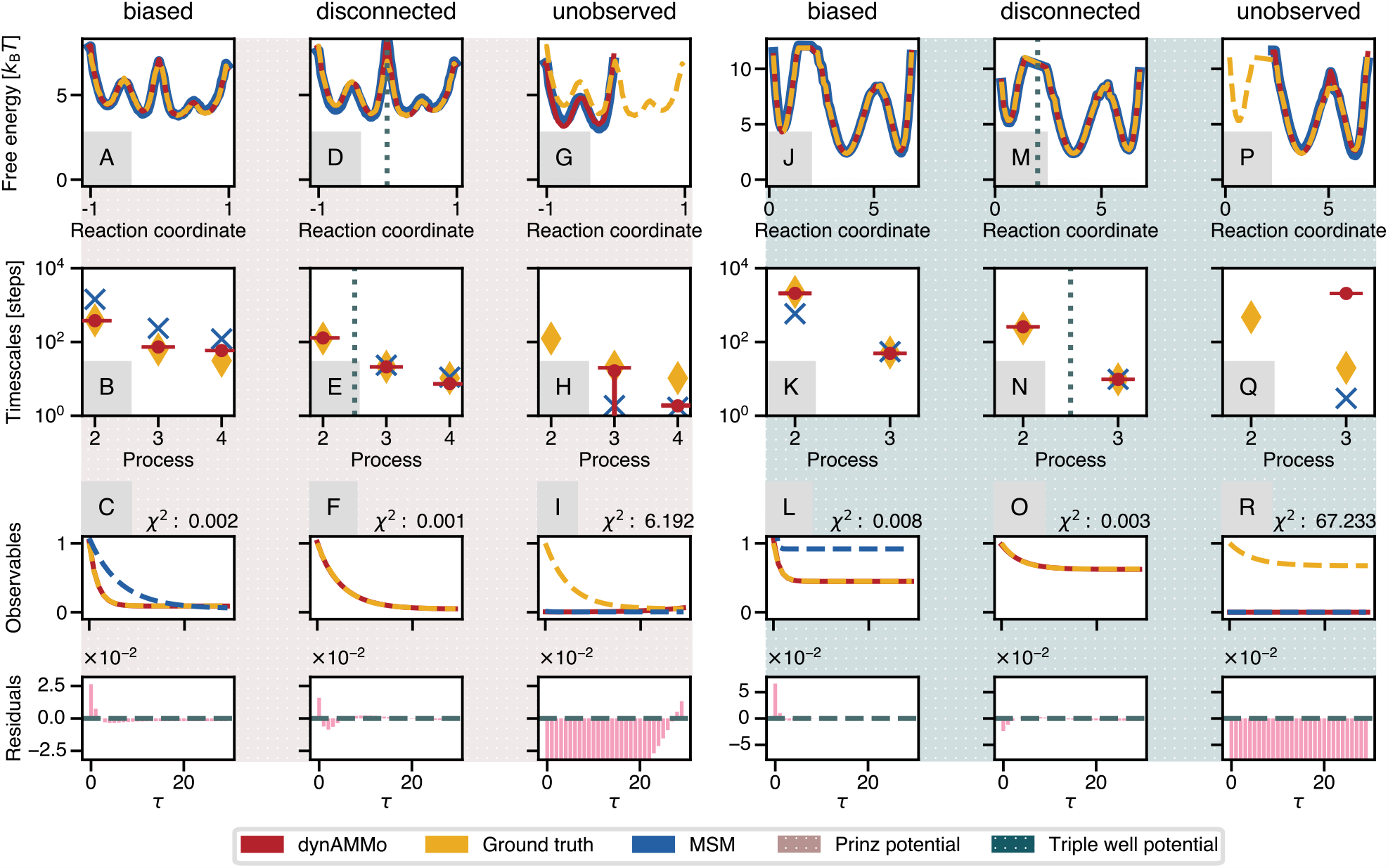
Overview of model system benchmark results. Three different scenarios using two model systems, the Prinz potential (brown background) and the three-well potential (teal background) are shown. The three different scenarios include biased simulations (A–C, J–L), disconnected trajectories (D–F, M–O), and unobserved or missing states (G–I, P–R). Each of the scenarios show the free energy potential, timescales of exchange, and observable plots. *χ*^2^ values and residuals between the model predictions and the ground truth data are also shown for the observable plots. The model results are shown in red, the ‘ground truth’ experimental data in yellow, and the naive MSM predictions in blue. ∆G values of the slowest transition for the different systems are given in the Supporting Information (Table S4).

We will first examine the scenario where we have biased simulations, which we compare with the experimental data derived from a “ground truth” model. In this case, all states were reversibly sampled in the simulations. However, due to, for example, force field inaccuracies and finite sampling, the timescales of exchange between the processes and the thermodynamics of the system do not correspond to the “true” ensemble. By investigating the free energy profiles of the two systems (Fig. 2A/J), we see that the populations of the MSM (blue) match the ground truth well (yellow). Consequently, we would not expect reweighing considering only the stationary observables, as is done in AMMs, will not have a big effect on the prediction of the dynamic observables since the timescales show significant discrepancies (Fig. 2B/K) between the MSM and the ground truth. Using dynAMMo, we can perfectly match the slowest timescales (Fig. 2B/K). We find a similar mismatch between the MSMs estimated on the biased data and the “ground truth data” (Fig. 2C/L). Since our observable function inherently informs about the slowest process we find that, dynAMMo does not substantially modify the timescale of the faster processes (Fig. 2B). This implies that the model does not introduce unnecessary bias into the estimation if it is not reflected in the observable. As opposed to the Prinz potential, we find that the predicted kinetics in the model trained on biased data from the three-well potential (Fig. 2, panels J–L) are accelerated compared to the ground truth (Fig. 2K). This mismatch in timescales manifests itself as poor agreement with the observable time correlation functions (Fig. 2L) between the ground truth (yellow) and the naive MSM (blue). Integrating the correlation function data and the biased simulation data with dynAMMo, we are in agreement with the ground-truth data and match the underlying rates.

### Disconnected simulations can be combined to a single Markov model using dynamic constraints

Many systems are characterized by timescales which are impractical to sample with statistical confidence. However, we may have access to multiple experimental structures of each of the states in isolation, some of which we can sample transitions between, others which are infeasible to sample. An example is the bovine pancreatic trypsin inihibitor (BPTI), for which numerous studies have reported slow millisecond timescale dynamics [94–97], and a millisecond long MD simulation only sampled the suspected slow transition once [16]. In many other cases, sampling such a transition in an unbiased fashion remains impractical.

To test such a scenario, we designed two experiments where we discard the transition counts of the slowest transition (Fig. 2D–E/M–N, gray dotted line) and split the trajectory in two. We build two Markov state models corresponding to the, now, disconnected subregions of the state space (Supporting Information, ‘Materials and Methods’). After reweighing the populations using the stationary AMM procedure [88], we perfectly match the model (red) and the ground truth (yellow) populations (Fig. 2D/M). Despite not having prior information on the slowest process, we can correctly identify the missing timescale using dynamic experimental observables (Fig. 2E/N). Our model bridges the two sides, even in the absence of observed transitions between them. This discovery is guided by the correlation function, i.e., the observable (Fig. 2F/O). The correlation function indirectly holds this information (Eq. S4), as a slower exchange process corresponds to a slower decay in the correlation function. Since we can match, both, the kinetics and the thermodynamics of the systems, we can also fit the observable prediction (Fig. 2F/O red, solid) to the ground truth (Fig. 2F/O yellow, dashed). Using experimental dynamic observables, we can thereby merge disconnected simulation statistics and estimate a single model which faithfully reproduces all the available data.

### dynAMMo does not overfit when relevant states are missing

Next, we consider the case where one or more states that contribute to a measurable experimental signal is missing from the MD simulation data. This situation is common in MD simulation studies as the simulation time is often insufficient to sample all the relevant states, and is an edge case related to the ‘disconnected’ situation discussed above. However, contrary to the previous case, we do not have all structural information to support a reliable prediction of the observable here. We therefore anticipate that dynAMMo is unable to yield a model perfectly fitting these data. To simulate this scenario, we discard transition counts from one state completely. Concretely, this procedure discards simulation data about states above 0 (Fig. 2G–I) in the Prinz potential and states below 2.5 in the asymmetric triple well potential (Fig. 2P–R). In this case, models built with dynAMMo cannot match (Fig. 2I/R) the data, which translates into missing timescales (Fig. 2H/Q). Mismatching predictions indicate that the model is failing, which suggests that one or more states that give rise to a measurable signal are missing.

### A mechanistic model of BPTI disulfide isomerization dynamics with dynAMMo

To test how our model performs in a realistic scenario, we turned to BPTI as a protein system. BPTI is a 58-residue protein whose dynamics has been extensively studied, both experimentally [95, 96, 98, 99] and computationally [16, 100]. BPTI is known to have micro-to millisecond conformational exchange [95, 96, 101], centered mainly on different isomerizations of the disulfide bond between Cys14 and Cys38. In addition, there is a 1-ms long MD trajectory available at a temperature of 300 K [16]. In this simulation, all known major conformations of BPTI are sampled, and the transitions between them show a distinct separation of timescales. Analysis of the trajectory shows conformational exchange in the fast microsecond regime [16, 88, 102], which is much faster than what the experimentally determined rates are suspected to reflect these processes. The discrepancy between the experiments and the simulation data makes BPTI an ideal test case for demonstrating dynAMMo as an avenue to reconcile the data.

By integrating simulation [16] and experimental CPMG data [94] from NMR spectroscopy with dynAMMo, we build a kinetic model of BPTI (Fig. 3). In line with previous analyses,[88, 103], we used time-lagged Independent Component Analysis (tICA) [104] to define a low dimensional space which we discretized into 384 states and aggregated into 4 metastable structural states. Consistent with previous analyses, the major structural substates display isomerization of the disulfide bridges between residues 14 and 38 (Fig. 3A). We show the most populated states colored purple and light blue (a total population of approximately 90 %), while the remaining population is shared by the green and orange states (Fig. 3A). The state connectivity is dense and rates vary across an order of magnitude (Fig. 3B), which we show with arrows of varying thickness between the states colored by identity and scaled by their relative populations. The slowest rates correspond to the transitions to the two minor states (Supporting Information, Fig. S6a) and we expect to occur in the low millisecond regime. The implied timescales computed from our dynAMMo model are systematically shifted compared to those of the naive Markov state model that only takes simulation statistics into account (Fig. 3C). For the MSM, the slowest processes are barely on the order of hundreds of microseconds (dark blue crosses) [91]. On the other hand, the slowest implied timescale estimated by dynAMMo is on the order of magnitude of approximately 3.2 ms. This timescale matches well with those estimated from experimental data at the same temperature using a two-state fit, where Millet et al. determined a chemical exchange of the order of 2 − 3 ms [94].

**Figure 3:**
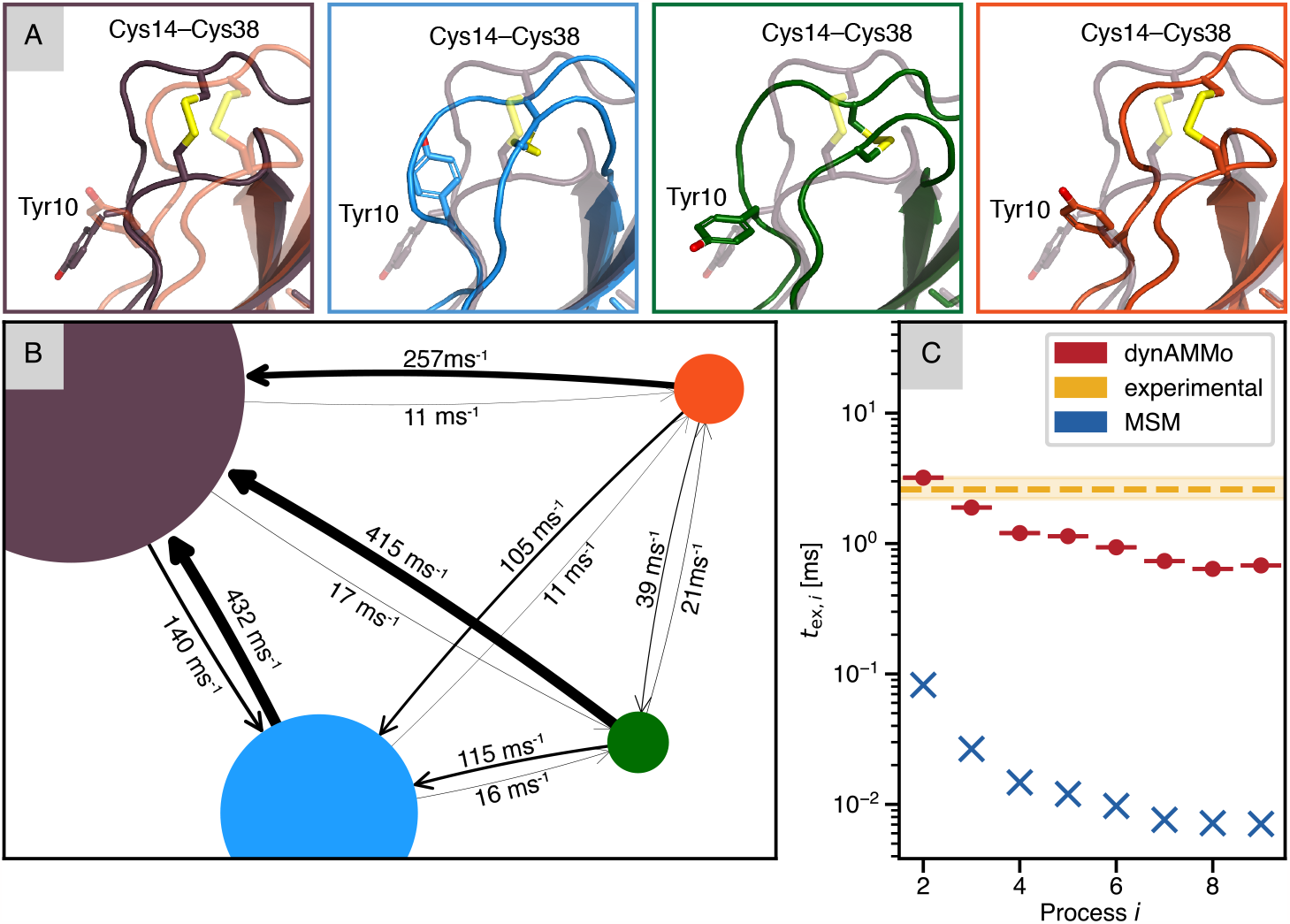
Integrating BPTI simulations and CPMG NMR data to build a quantitative kinetic model with dynAMMo. (A) Representative structures of the major BPTI states with the structural characteristics highlighted. Aromatics are shown for orientation. (B) Kinetic network between macrostates. The colors of the nodes correspond to the structural representatives shown above. The size of the nodes and the arrow widths are proportional to the size of the populations as well as the reaction rate, respectively. Rates are shown above / below the arrows in ms^−1^. (C) Timescales of exchange as a function of the slowest processes. dynAMMo is shown in red and the MSM from the simulation data is colored blue. The time constant of the experimentally determined exchange is shown in yellow with the standard deviation shown as shaded area.

In Fig. 4 we show representative examples of some key observables. The plots show CPMG relaxation dispersion curves, which measure the effect of chemical exchange on ^15^N^H^ spins. The chemical shift predictions that were used as observables for the backbone amides were obtained by the PPM algorithm [105]. The experimental CPMG data are shown in yellow, whereas the predictions of the model are shown in red and the predictions were scaled according to the values reported in the Supporting Information (Fig. S7). All observables show an excellent overall agreement with the data, suggesting that the underlying model is capable of explaining the data in a meaningful way (see Supporting Information, Fig. S8 and S9). We note that all relevant residues involved in the exchange display a relatively strong dispersion, which we are able to perfectly match using dynAMMo. This observation strengthens the argument that the conformations sampled in the MD simulation constitute the relevant configurations needed to explain the experimental data. We stress that dynAMMo uses all observables to fit one global kinetic model. Therefore, the predictions of the relaxation dispersion curves for the backbone nitrogens differ only by the observable used for each residue. Here, we demonstrate how dynAMMo can be used to combine experimental NMR relaxation data with simulation data from molecular dynamics simulations. Therefore, we obtain a detailed mechanistic explanation of how the different metastable states observed in the MD trajectory contribute to the experimentally probed chemical exchange.

**Figure 4:**
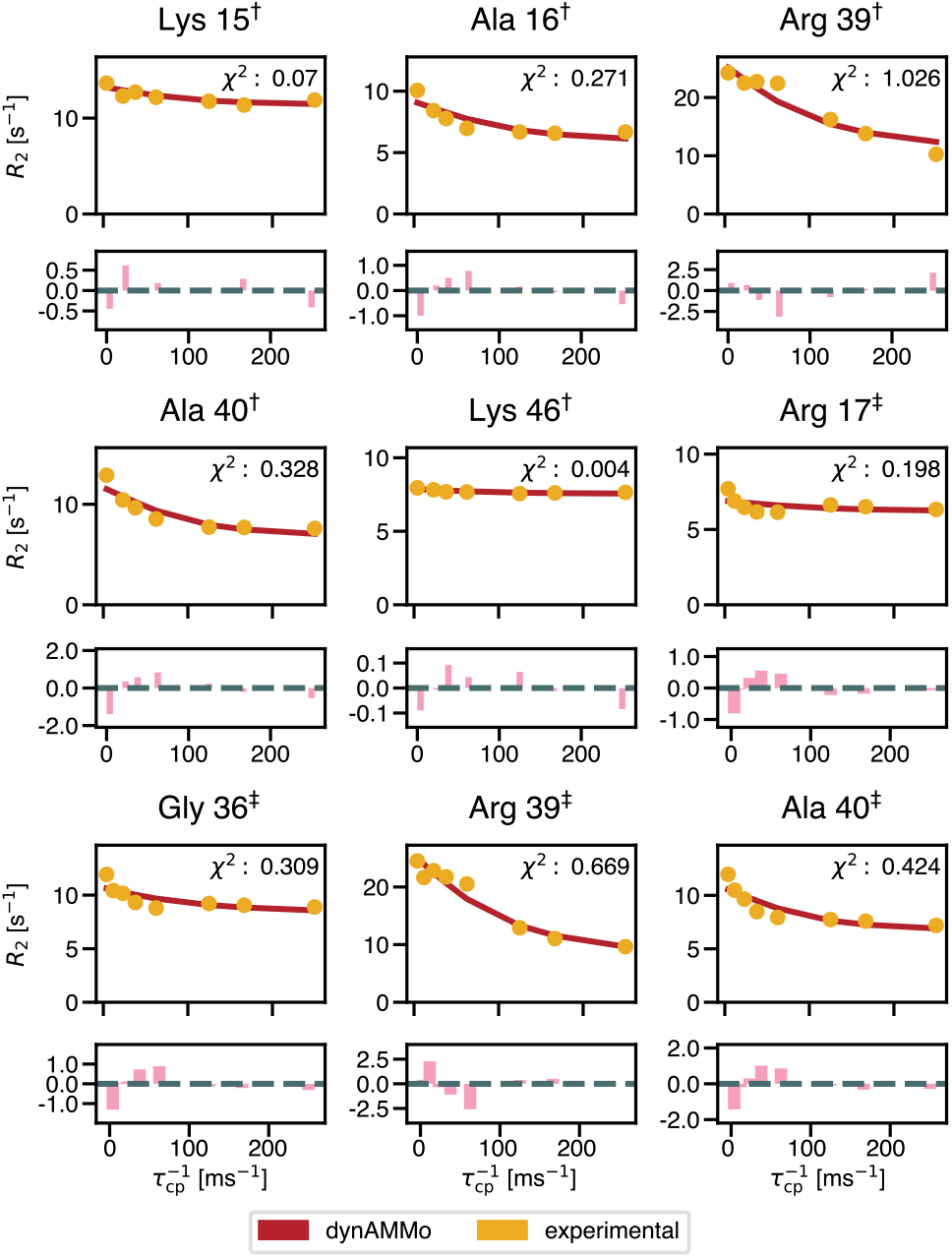
CPMG relaxation dispersion data of BPTI ^15^N^H^ spins. Nine representative examples of CPMG plots are shown. The fitted model (red, solid) is in high agreement with the experimental data (yellow circles). *χ*^2^ values between the prediction and the experiments are shown for each subplot. The dagger refers to the dataset that has been recorded at a Larmor frequency of 600 MHz. Conversely, the double dagger refers to the dataset recorded at 500 MHz [94]. Residuals between the experiments and predictions are shown in pink.

### Quantitative molecular kinetics from disconnected simulation statistics with dynAMMo

Above we saw how dynAMMo could recover the correct kinetics on a controlled test system. To evaluate whether dynAMMo generalizes to more complex protein systems, we establish a similar benchmark, systematically removing simulation statistics that connect the major and minor populated states of BPTI. For this case, we similarly find that dynAMMo can quantitatively recover the exchange rates between the disconnected states (Fig. 5A), and recover the implied timescales accurately (Fig. 5B), despite the minor discrepancies between the timescales in the connected and disconnected case (Fig 3 and 5). The discrepancies observed, although noticeable, are on the same order of magnitude. We are to expect these due to a combination of limited data and data uncertainty. The MSM in this case is missing the slowest process (Fig. 5A, dashed cross), however, dynAMMo can recover this process and quantitatively predict the timescale. We show a detailed analysis of this scenario in the Supporting Information (Fig. S6b).

**Figure 5:**
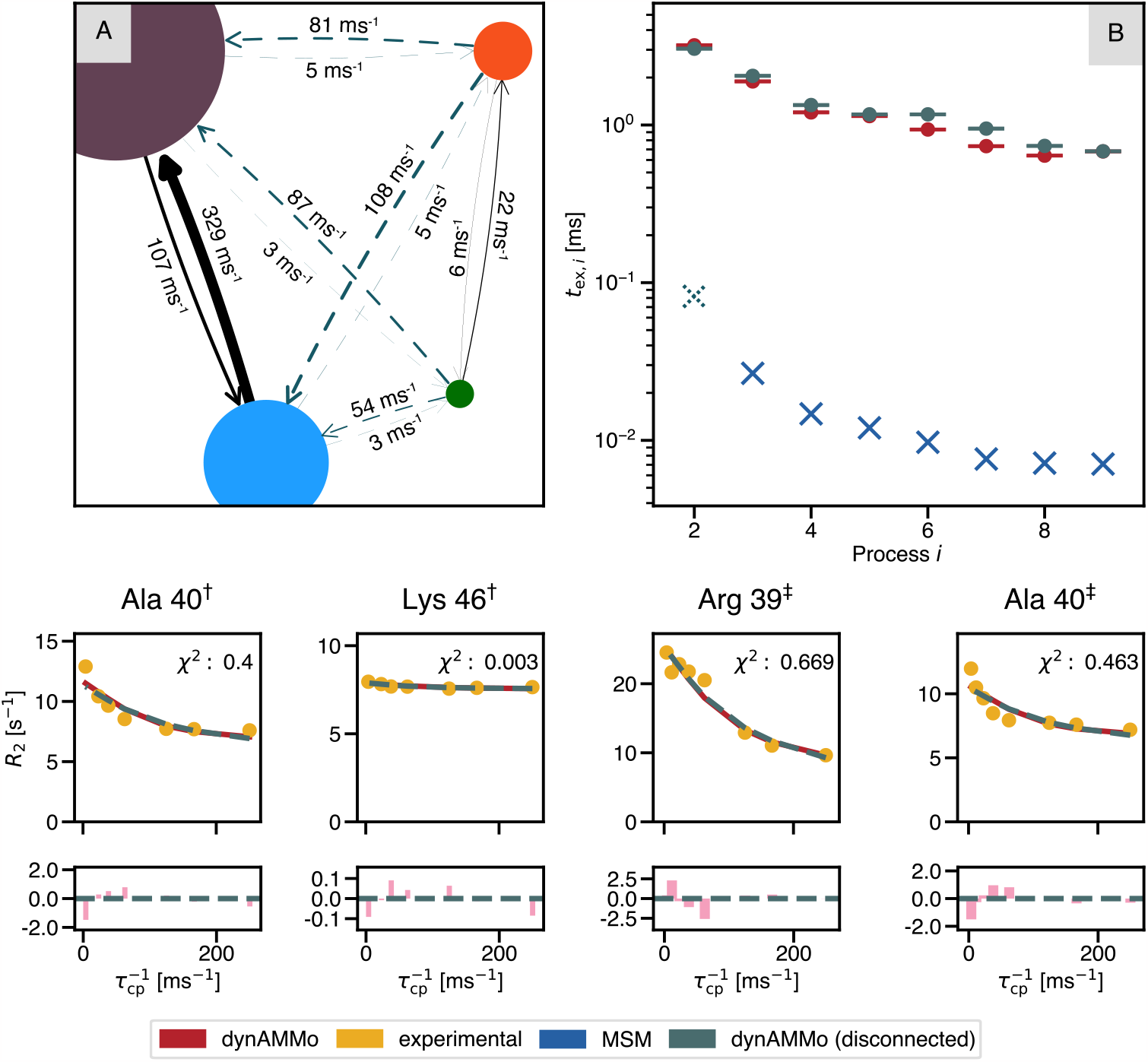
Overview of simulating disconnected case using BPTI simulations and CPMG data. (A) Kinetic network between the disconnected states. All transitions between the purple/light blue and orange/green clusters were removed (indicated with dashed gray arrows) and two Markov state models were built using only the within-states trajectory data. The colors of the states correspond to the clusters shown in 3A. (B) Implied timescale plot of the disconnected model (gray), the connected model (red), and the MSM (blue) for comparison. The presumed timescale of exchange that was removed in this scenario is indicated as a dashed cross. Lower panel: Four representative examples of CPMG relaxation dispersion predictions of selected backbone nitrogens. The predictions of the disconnected case (dashed gray) are plotted together with the predictions of the connected scenario (red) for comparison. *χ*^2^ values are shown with respect to the predictions of the disconnected case and the experimental data (yellow).

## Conclusion

Here we have introduced dynamic Augmented Markov Models (dynAMMo), a new method to improve the accuracy of mechanistic biomolecular models by incorporating dynamic experimental measurements to correct for biases in the kinetics and thermodynamics of MD simulations. However, most intriguingly, dynAMMo also allows us to build quantitatively predicted model of molecular kinetics even in the absence of simulation statistics on (slow) conformational transitions. We show the performance of dynAMMo across two well-controlled benchmark systems and later deploy it to two realistic scenarios using data from molecular simulations and NMR spectroscopy on the protein BPTI. It is essential to highlight that while dynAMMo offers significant advancements, the robustness of the model still depends on the quality of the initial MD and experimental data. In cases where MD simulations inadvertently do not sample certain rare events or states, we can only build a (useful) dynamic Augmented Markov Model if all states are known but only some transitions are missing (“disconnected” model case). The model fails if one or more important state are missing, offering important insights into the conformational space of the molecule. This observation reinforces the need for comprehensively sampling the relevant conformational states, either through simulations or with experimental techniques. Further, since dynAMMo balances experimental and simulation data through the principle of maximum entropy, we approach a compromise from an information theoretical perspective. Therefore, although multiple models could potentially explain the data, we identify the one that requires minimal perturbation from the simulation data to align with the available experimental evidence. As such, dynAMMo opens up the possibility of salvaging sparsely sampled simulation data sampled using biased force fields and repurpose them to build quantitatively predictive models for structural biology.

## Materials and Methods

The estimator is implemented in Python and uses PyTorch [106] and deeptime [107] as the main analysis and modeling tools. The estimation procedure and theory details are provided in Supporting Information, ‘Theory’. The code will be made available on https://github.com/olsson-group/dynAMMo. All figures showing molecular structures were made using PyMol [108]. All plots were generated using Matplotlib [109]. Additional results, such as the slowest estimated eigenvectors, loss function, and stationary distribution are reported in the Supporting Information for all model systems (Fig. S4 and S5) and BPTI (Fig. S6), respectively.

## Benchmark model systems

The deeptime implementation of the four-state Prinz potential and the three-well potential datasets was used to simulate the two benchmark systems [107]. The parameters used to simulate the trajectories are reported in the Supporting Information (table S1). The estimation of the dynamic Augmented Markov Models were carried out as outlined in *Results and Discussion*. Each scenario uses different parametrizations of the potential and a table with an overview is listed in the Supporting Information (Table S2). Chapman-Kolmogorov tests have been performed on all MSMs used in this study (Supporting Information Fig. S1–S2).

## BPTI

The estimation and analysis of the BPTI dynAMMo were conducted as described in *Results and Discussion*. Chapman-Kolmogorov tests were conducted on the MSMs used here to ensure validity of the models Supporting Information (Fig. S3). The estimation parameters of the two scenarios are listed in the Supporting Information (Table S3). Further details are provided in Supporting Information, ‘Materials and Methods.’

## Acknowledgements

The authors would like to thank D. E. Shaw Research for sharing the BPTI simulation and Arthur G. Palmer III, for sharing the raw NMR relaxation dispersion data. CK thanks Shanawaz Ahmed for fruitful discussions and for sharing a preliminary implementation of the Cayley transform for the eigenvector estimation. This work was partially supported by the Wallenberg AI, Autonomous Systems and Software Program (WASP) funded by the Knut and Alice Wallenberg Foundation.

## Author Contributions

C.K. and S.O. conceptualized, designed, and performed research as well as wrote the manuscript; C.K. analyzed and visualized the data and performed statistical analysis; S.O. provided supervision, project administration and funding acquisition.

