## Supplemental Material for "Rescuing Off-Equilibrium Simulation Data through Dynamic Experimental Data with dynAMMo"

#### This PDF file includes:

Supporting text
Figs. S1 to S9
Tables S1 to S4
SI References

### Supporting Information Text

### 1. SI Theory

Markov state models (MSMs) are a powerful framework to characterize the dynamics of biomolecular systems (1–3). With dynamic Augmented Markov Models (dynAMMo), we leverage the expressivity of standard Markov models and combine them with dynamic experimental observables to obtain a more comprehensive understanding of biomolecular systems. Markov models are a master-equation framework, describing the system's dynamics solely with the transition matrix $\mathbf{T}$ at lag time $\tau$:

$$\mathbf{T}(\tau) = \mathbf{L}\mathbf{\Lambda}(\tau)\mathbf{R}, \quad \text{[S1]}$$

where $\mathbf{L}$ and $\mathbf{R}$ are the left and right eigenvectors, respectively, and $\mathbf{\Lambda}$ is the diagonal matrix of the eigenvalues. They constitute the spectral components of $\mathbf{T}$. $\mathbf{L}$ is defined as

$$\mathbf{L} = (\mathbf{\Pi}\mathbf{R})^\top \quad \text{[S2]}$$

with $\mathbf{\Pi} = \mathrm{diag}(\boldsymbol{\pi})$.

With dynamic Augmented Markov Models, we maximize the likelihood of $\hat{\mathbf{T}}$ using constrained optimization and maximum entropy principles guided by the experimental observations by modifying the aforementioned spectral components of $\hat{\mathbf{T}}$.

#### A. Estimation of dynamic Augmented Markov Models.

***A.1. Experimental loss.*** At the core of our approach, we want to match the predictions of our model $\mathbf{o}^{\text{pred}}$ to the experimental data $\mathbf{o}^{\text{exp}}$. Matching is achieved by calculating the mean squared error of the two quantities multiplied by a weight factor $\mathbf{W}$:

$$\mathcal{L}^{\text{experimental}} = \sum_{l,k} \sigma(w_{l,k})(o_{l,k}^{\text{pred, dynamic'}} - o_{l,k}^{\text{exp, dynamic}})^2 + \sigma(w_{l,k})^{-1} - 1. \quad \text{[S3]}$$

Here, $o^{\text{pred, dynamic'}} = m \cdot o^{\text{pred, dynamic}} + b$, where $m$ and $b$ are two positive constants. They are necessary for example for predicting relaxation dispersion experiments, where $b = R_{2,\text{intrinsic}}$ and $m$ is a correction factor to recover the absolute scale (4). Note that $m$ can be quite large due to a range of factors, including the discretization, and subsequent averaging of the observable within the Markov states. Underestimation of the variance in the experimental observable is best quantified by comparing model predictions at zero lag-time (eq. S1) to the empirical sample variance. The weights are mapped to the interval $[0, 1]$ by the sigmoid function defined as $\sigma(x) = \frac{1}{1+e^{-x}}$. The loss is augmented by the prior $\sigma(\mathbf{W})^{-1} - 1$ that penalizes lower weights, thus ensuring that informative data points are given high importance.

***A.2. Connection of MSMs to experiments.*** $\mathbf{o}^{\text{pred}}$ is calculated using equation 1. More specifically, equation 1 is equivalent to the multiexponential sum of all processes $i$:

$$o^{\text{dyn}}(k) = \sum_{i}^{m} c_i \exp\left(-\frac{k\tau}{t_i^{\text{ex}}}\right) \quad \text{[S4]}$$

with

$$t_i^{\text{ex}} = -\frac{\tau}{\log(|\lambda_i|)} \quad \text{and} \quad c_i = (\mathbf{a}^\top \mathbf{L}_i)^2. \quad \text{[S5]}$$

Equations 1 and S5 are the basis for calculating quantities, such as the transverse relaxation rate $R_2^{\text{ex}}$:

$$R_2^{\text{ex}} = (2\pi\nu_0)^2 \sum_{i=2}^{n} c_i t_i^{\text{ex}}, \quad \text{[S6]}$$

the rotating-frame relaxation rate $R_{1\rho}^{\text{ex}}$:

$$R_{1\rho}^{\text{ex}}(\nu_1) = (2\pi\nu_0)^2 \left(\sum_{i=2}^{n} c_i \frac{t_i^{\text{ex}}}{1 + (t_i^{\text{ex}}\nu_1)^2}\right), \quad \text{[S7]}$$

as well as Carr-Purcell-Meiboom-Gill (CPMG) relaxation rate (4):

$$R_{\text{CPMG}}^{\text{ex}}(\nu_{\text{CP}}) = (2\pi\nu_0)^2 \left(\sum_{i=2}^{n} c_i t_i^{\text{ex}} \left(1 - \frac{t_i^{\text{ex}}}{\tau_{\text{CP}}} \tanh\frac{\tau_{\text{CP}}}{t_i^{\text{ex}}}\right)\right) \quad \text{[S8]}$$

They allow us to directly compare observables derived from Markov models with their experimental counterparts $\mathbf{o}^{\text{pred}}$.

**A.3. Lagrange Loss.** Additional constraints on the model are enforced through Lagrange multipliers. Those include ergodicity ($\xi$), reversibility ($\varphi$), stochasticity ($\zeta$), and validity ($\chi$). The constraints correspond to ensuring a connected state-space, detailed balance, and a valid row-stochastic matrix. The total '*Lagrange*' loss consequently takes the form

$$\mathcal{L}^{\text{Lagrange}} = \sum_i \xi_i(\max(1, \lambda_i) - 1) \quad \text{[S9]}$$

$$+ \sum_{i,j} \varphi_{ij}(\pi_i p_{ij} - \pi_j p_{ji}) \quad \text{[S10]}$$

$$+ \sum_i \zeta_i(\sum_j p_{ij} - 1) \quad \text{[S11]}$$

$$+ \sum_{i,j} \sigma(\chi_{i,j})(-\min(0, T_{i,j})) + \sigma(\chi_{i,j})^{-1} - 1. \quad \text{[S12]}$$

Notice that the ergodicity and validity constraints, $\xi$ and $\chi$, are only active if the solution is outside the feasible region. That is, the constraints are violated.

**A.4. Maximum Entropy contribution.** The loss is further complemented with a Kullback-Leibler divergence between the stationary distribution of the reweighted model and the MSM is minimized, from Augmented Markov models (5),

$$\mathcal{L}^{\text{NLL}} = -\sum_i \hat{\pi}_i \ln\left(\frac{\hat{\pi}_i}{\pi_i}\right). \quad \text{[S13]}$$

**A.5. Total loss function.** The total loss function thus consists of a weighted sum of the aforementioned loss terms:

$$\mathcal{L}^{\text{total}} = \sigma(\alpha)\mathcal{L}^{\text{NLL}} + \beta\mathcal{L}^{\text{experimental}} + \mathcal{L}^{\text{Lagrange}}, \quad \text{[S14]}$$

where $\alpha$ and $\beta$ are hyperparameters.

**A.6. Update of eigenvectors.** Since our dynamic experimental observables are functions of the eigenvectors and eigenvalues of the estimated transition matrix, in practice we estimate the dynAMMo model by parameterizing its spectral components, e.g. its eigenvalues and eigenvectors. However, each right eigenvector $\mathbf{r}_i$ is orthonormal with respect to all other right eigenvectors $\mathbf{R}_{\neg i}$. In effect, we update the eigenvalues and eigenvectors directly when optimizing the loss (eq. S14), yet, we need do so in a manner which satisfies the constraints of these parameter: orthonormality and $|\lambda_i| \leq 1$. Optimization under such constraints happen on the Stiefel manifold(6), and is common in many machine learning problems (7–9). Our aim is to minimize to estimate the right eigenvectors, $\hat{\mathbf{R}}$, using the following expression:

$$\min_{\hat{\mathbf{R}} \in \mathbb{R}^{n\times n}} \mathcal{L}^{\text{total}}(\hat{\mathbf{R}}), \text{ such that } \hat{\mathbf{R}} \in \text{St}(n,n), \quad \text{[S15]}$$

where

$$\text{St}(n,n) = \left\{\mathbf{X} \in \mathbb{R}^{n\times n} : \mathbf{X}^\top \mathbf{X} = \mathbf{I}_n\right\} \quad \text{[S16]}$$

is the Stiefel manifold and $\mathbf{I}_n$ is the identity matrix in $n$-dimensions. Moving along this manifold is non-trivial, as the constraints are non-convex and numerically expensive, as they have to be preserved during every step of the iteration. Here, we employ the Cayley transform to update the gradients of the right eigenvectors (10) during estimation. For a $\hat{\mathbf{R}}$ and its gradient $\mathbf{G} \coloneqq \nabla_{\hat{\mathbf{R}}}\mathcal{L}^{\text{total}}$, we update

$$\hat{\mathbf{R}}' = \hat{\mathbf{R}} - \frac{\eta}{2}\mathbf{A}\left(\hat{\mathbf{R}} + \left(I + \frac{\eta}{2}\mathbf{A}\right)^{-1}\left(I + \frac{\eta}{2}\mathbf{A}\right)\hat{\mathbf{R}}\right), \quad \text{[S17]}$$

where $\mathbf{A} \coloneqq \mathbf{G}\hat{\mathbf{R}}^\top - \hat{\mathbf{R}}\mathbf{G}^\top$ and $\eta$ is the learning rate. We compute $\mathbf{G}$ using autograd functionality of PyTorch.

**B. Algorithm.** Here, we describe the initialization procedure (1)and the main estimation algorithm (2) for dynamic Augmented Markov models. For the connected case, e.g. the number of connected MSMs, $n = 1$, the initialization is equivalent to standard Markov state modelling (1, lines 16-18). For the disconnected case, i.e., $n > 1$, the user can choose between initializing the model using the count matrix or the transition matrix (line 3). In both cases, the matrices are assembled into a block matrix, where the off-diagonal values are either low transition probabilities (line 7) or low transition counts (line 11). it is important to note that for estimation, only the unmodified simulation data (counts) are considered. Once the model is initialized, the parameters are estimated according to algorithm 2.

The eigenvectors, eigenvalues and stationary distributions are all treated as parameters and are initialized from $\mathbf{T}^{\text{init}}$ (algorithm 2, line 2). The observable function $f$ and the lag times $k$ are chosen depending on the type of experimental data $\mathbf{o}^{\text{exp}}$ (line 2). The scaling parameters $\alpha$ and $\beta$ as well as the learning rate $\eta$ are hyperparameters and therefore have to be chosen by the user, whereas the coefficients of the affine data transform for the dynamic observables are treated as parameters but can be initialized by the user (line 1). Finally, the Lagrange multipliers are initialized using samples from a standard

**Algorithm 1** dynamic Augment Markov Model initialization

Initialize set of $n$ transition and count matrices, $\mathcal{T} = \{\mathbf{T}_1, \mathbf{T}_2, ...\mathbf{T}_n\}$ and $\mathcal{C} = \{\mathbf{C}_1, \mathbf{C}_2, ...\mathbf{C}_n\}$, as well as observables mapped onto Markov states $\mathcal{A} = \{\mathbf{a}_1, \mathbf{a}_2, ...\mathbf{a}_n\}$
Set `count_matrix_initialization` $\leftarrow$ True/False
**if** $n > 1$ **then**
    **if** `count_matrix_initialization` is True **then**
        $\mathbf{T}^{\text{init}} \leftarrow \text{block\_diag}(\mathcal{T})$
        $i \leftarrow$ off-diagonal indices of $\mathbf{T}$
        $\mathbf{T}^{\text{init}}[i] \leftarrow \epsilon,$ default: $10^{-10}$ ▷ Fill off-diagonals of block matrix with low transition probability
    **else**
        $\mathbf{C}^{\text{init}} \leftarrow \text{block\_diag}(\mathcal{C}) \times 10^3$ ▷ Multiply with constant factor
        $i \leftarrow$ off-diagonal indices of $\mathbf{C}$
        $\mathbf{C}^{\text{init}}[i] \leftarrow \epsilon,$ default: 1 ▷ Fill off-diagonals of block matrix with low counts
        $\mathbf{T}^{\text{init}} \leftarrow \text{MarkovStateModel}(\mathbf{C}^{\text{init}})$ ▷ Estimation according to (11)
    $\mathbf{C} \leftarrow \text{block\_diag}(\mathcal{C})$
    $\mathbf{a} \leftarrow \text{concatenate}(\mathcal{A})$
**else**
    $\mathbf{T}^{\text{init}} \leftarrow \mathbf{T}_1$
    $\mathbf{C} \leftarrow \mathbf{C}_1$
    $\mathbf{a} \leftarrow \mathbf{a}_1$
$\mathbf{M} \leftarrow \text{DynamicAugmentedMarkovModel}(\mathbf{T}^{\text{init}}, \mathbf{C}, \mathbf{a})$

normal distribution. If the set of experiments contains stationary observables, the stationary distribution is estimated via the original Augmented Markov Model proposed by (5) (line 11). It is, however, possible to omit providing stationary experiments (for example if they are not available), and only use dynamic observables. In this case, the stationary distribution is treated as a parameter that is estimated via gradient descent. In the main estimation algorithm, we update our predictions of dynamic observables based on the current set of parameters (line 19). The transition matrix is then updated and all losses are calculated according to equations S3-S14 (lines 21-24). A backward pass is performed and the eigenvectors are updated as described in section A.6. The other parameters are updated via gradient descent (12). Convergence is reached when the median of the loss "acceleration" (change of change of the total loss) is below a certain threshold or a certain number of iterations.

**Algorithm 2** dynamic Augment Markov Model

1: Initialize $\hat{\mathbf{R}}, \boldsymbol{\lambda}, \hat{\boldsymbol{\pi}} \leftarrow \mathbf{T}^{\text{init}}$ ▷ initialization via algorithm 1
2: Initialize experimental data $\mathbf{o}^{\text{exp}}$, independent variable $k$, and observable function $f$
3: Initialize affine coefficients $a, b$, scaling parameters $\alpha, \beta$ and learning rate $\eta$
4: Initialize $\hat{\boldsymbol{\theta}} \leftarrow \{\boldsymbol{\lambda}, \hat{\boldsymbol{\pi}}, \hat{\mathbf{R}}, \text{ Lagrange multipliers}, a, b, \alpha, \beta, \gamma\}$
5: Set `converged` $\leftarrow$ False
6:
7: **if** StationaryExperiment $\in$ experiments **then**
8:     **repeat**
9:         **for** $\mathbf{o}^{\text{exp, stationary}} \in$ experiments **do**
10:             $\mathbf{o}^{\text{pred, stationary}} \leftarrow \text{StationaryExperiment}(\mathbf{a}, \hat{\boldsymbol{\pi}})$
11:         $\hat{\boldsymbol{\pi}} \leftarrow \text{stationaryAMM}(\mathbf{o}^{\text{exp, stationary}}, \mathbf{o}^{\text{pred, stationary}}, \hat{\boldsymbol{\pi}})$ ▷ implementation according to (5)
12:         $\hat{\boldsymbol{\theta}} = \hat{\boldsymbol{\theta}} \setminus \{\hat{\boldsymbol{\pi}}\}$
13:     **until** converged
14: **else**
15:     $\hat{\boldsymbol{\pi}}$ is updated via gradient descent
16:
17: **while** not `converged` **do**
18:     **for** $\mathbf{o}^{\text{exp, dynamic}} \in$ experiments **do**
19:         $\mathbf{o}^{\text{pred, dynamic}} \leftarrow \text{DynamicExperiment}(k, f, \mathbf{a}, \hat{\boldsymbol{\pi}}, \boldsymbol{\lambda}, \hat{\mathbf{R}}, m, b)$
20:     $\hat{\mathbf{T}} = \text{update}(\mathbf{C}, \hat{\boldsymbol{\theta}})$
21:     Calculate $\mathcal{L}^{\text{experimental}}(\mathbf{o}^{\text{exp, dynamic}}, \mathbf{o}^{\text{pred, dynamic}}, a, b)$ ▷ equation S3
22:     Calculate $\mathcal{L}^{\text{Lagrange}}(\hat{\boldsymbol{\theta}})$ ▷ equation S12
23:     Calculate $\mathcal{L}^{\text{NLL}}(\mathbf{C}, \hat{\mathbf{T}})$ ▷ equation S13
24:     Calculate $\mathcal{L}^{\text{total}}(\mathcal{L}^{\text{Lagrange}}, \mathcal{L}^{\text{NLL}}, \alpha, \beta)$ ▷ equation S14
25:     Backpropagate
26:     Project gradients of $\hat{\mathbf{R}}$ onto Stiefel manifold $\Delta\hat{\mathbf{R}} = \hat{\mathbf{R}} - \eta_{\text{R}} \frac{\partial \mathcal{L}}{\partial \hat{\mathbf{R}}}$ ▷ equation S17
27:     Update other parameters $(\hat{\boldsymbol{\theta}}' = \hat{\boldsymbol{\theta}} \setminus \{\hat{\mathbf{R}}\})$: $\Delta\hat{\boldsymbol{\theta}}' = \hat{\boldsymbol{\theta}}' - \eta \frac{\partial \mathcal{L}}{\partial \hat{\boldsymbol{\theta}}'}$
28:     **if** $\text{median}(\Delta\Delta\mathcal{L}^{\text{total}}) < \varepsilon$ or `n_iterations` $\geq$ `n_total` **then**
29:         `converged` = True
30:         $\mathbf{M} \leftarrow \text{DynamicAugmentedMarkovModel}(\hat{\mathbf{T}}, \mathbf{C}, \mathbf{a})$
31:     **else**
32:         continue

### SI Materials and Methods

**C. dynAMMo and MSM estimation of benchmark systems.** Trajectories were generated using the Prinz potential (1) and the triple-well potential using the deeptime library (11) at different temperatures for the different scenarios (see table S1). The trajectories were discretized into 100 and 69 states, respectively. MSMs were estimated using deeptime and the lag times described in table S2. The validity of the estimated Markov models was verified using Chapman-Kolmogorov (CK) tests (shown in figure S1 and S2. dynAMMo was then estimated using the parameters listed in table S2.

**D. dynAMMo and MSM estimation of BPTI.** All backbone torsions as well as the side chain torsions of residues 13-14 and residues 37-38 were used as features for time-lagged Independent Component Analysis (tICA) estimation with a lag time of 500ns using a 1-ms trajectory. The simulations were carried out as reported (13). The largest eight independent components were used for K-Means clustering using 384 centres. The discretized trajectory was used for count matrix estimation using a lag time of 1µs. MSM estimation was carried out using the count matrix followed by self-consistency checks (implied timescale plots and Chapman-Kolmogorov tests, see figure S3). The dynamic Augmented Markov models were built using the parameters shown in table S3. The initial values for the correction factor $m$ and the intrinsic transverse relaxation rate $b = R_{2,\mathrm{intrinsic}}$ (see equation S3) for predicting the CPMG observables were calculated as $m_{i,\mathrm{init}} = o_i^{\mathrm{exp}}[0]/\mathrm{std}(o_i^{\mathrm{exp}})$ and $b_{i,\mathrm{init}} = o_i^{\mathrm{exp}}[-1]$ for every observable $i$. Here, [0] and [-1] refer to the first and last data point of each observable $i$ in the series, respectively. The estimated parameters are shown in figure S7.

| | | Prinz potential | | | Triple-well potential | | |
|---|---|---|---|---|---|---|---|
| | | biased | disconnected | missing | biased | disconnected | missing |
| Simulation | `n_steps` | 1.00E+03 | 5.00E+02 | 5.00E+02 | 2.00E+03 | 2.00E+03 | 2.00E+03 |
| | `step_size` | 1.00E-06 | 1.00E-05 | 1.00E-05 | 1.80E-02 | 1.80E-02 | 1.80E-02 |
| "Ground truth" | `n_steps` | 2.00E+03 | 5.00E+02 | 5.00E+02 | 1.90E+03 | 2.00E+03 | 2.00E+03 |
| | `step_size` | 1.00E-07 | 1.00E-05 | 1.00E-05 | 1.80E-03 | 1.80E-02 | 1.80E-02 |

**Table S1. Parameter overview for simulating the trajectories of the model systems.**

| | Prinz potential | | | Triple well potential | | |
|---|---|---|---|---|---|---|
| | connected | disconnected | unobserved | connected | disconnected | unobserved |
| `lag` | 15 | 5 | 5 | 30 | 8 | 10 |
| $\alpha$ (scaling factor) | -9 | -10 | -10 | -9 | -8 | -12 |
| $\beta$ (scaling factor) | 1.00E+03 | 1.00E+03 | 1.00E+03 | 1.00E+01 | 1.00E+03 | 1.00E+00 |
| $\eta$ (learning rate) | 1.00E-03 | 1.00E-03 | 1.00E-03 | 1.00E-02 | 1.00E-02 | 1.00E-06 |
| $\eta_{\mathrm{R}}$ (learning rate) | 1.00E-16 | 5.00E-15 | 1.00E-14 | 1.00E-13 | 1.00E-17 | 1.00E-14 |
| $\varepsilon$ (convergence threshold) | 2.00E-02 | 2.00E-02 | 2.00E-02 | 1.00E-02 | 3.00E-02 | 2.00E-02 |
| `n_eigvals` | 3 | 3 | 3 | 2 | 2 | 2 |

**Table S2. dynAMMo parameters for different benchmark scenarios.**

| | BPTI connected | BPTI disconnected |
|---|---|---|
| `lag` | 1µs | 1µs |
| `dt_traj` | 1.00E-06 | 1.00E-06 |
| $\alpha$ (scaling factor) | -15 | -15 |
| $\beta$ (scaling factor) | 1 | 1 |
| $\varepsilon$ (convergence threshold) | 0.02 | 0.02 |
| $\eta$ (learning rate) | 1.00E-04 | 1.00E-05 |
| $\eta_{\mathrm{R}}$ (learning rate) | 1.00E-20 | 1.00E-20 |
| `n_eigvals` | 8 | 8 |

**Table S3. dynAMMo parameters for BPTI connected and disconnected case.**

**E. Chapman-Kolmogorov tests.** Plots S1 and S2 show the estimated (from trajectory data) and predicted (from MSM) transition probabilities of the Prinz potential and triple-well potential MSMs. The lag times are shown in table S2. The metastable states were calculated using PCCA (11).

Figure S3 shows the CK tests of the BPTI MSMs used in this study. The lag times at which the predictions are made us shown in table S3.

### SI results

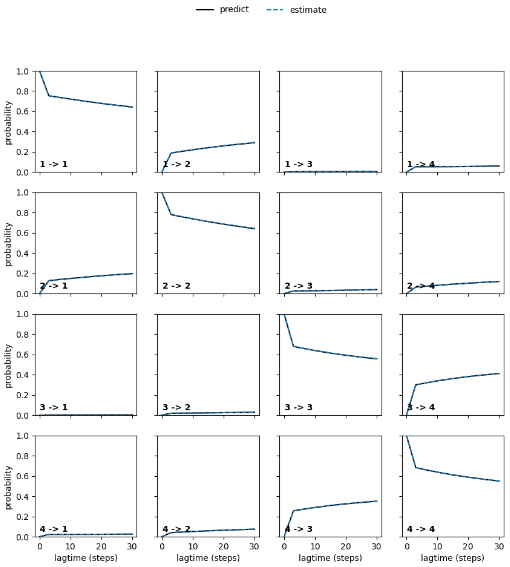


**(a)** Prinz potential (connected) MSM Chapman-Kolmogorov test.

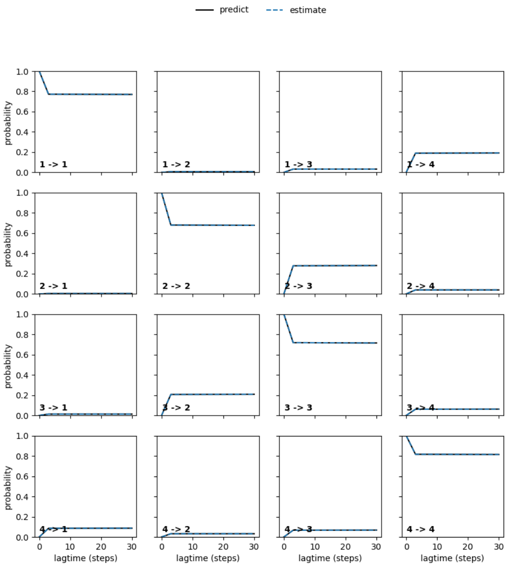


**(b)** Prinz potential (missing) MSM Chapman-Kolmogorov test.

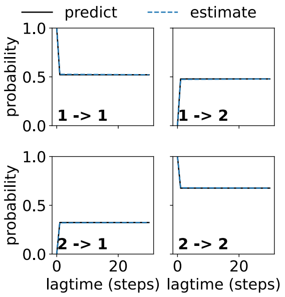


**(c)** Prinz potential (disconnected, 0-49 states) MSM Chapman-Kolmogorov test.

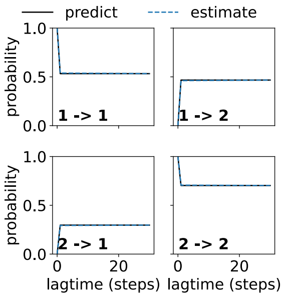


**(d)** Prinz potential (disconnected, 50-100 states) MSM Chapman-Kolmogorov test.

**Fig. S1.** Prinz potential potential CK tests of all MSMs: connected (a), missing state (b) and disconnected (c and d).

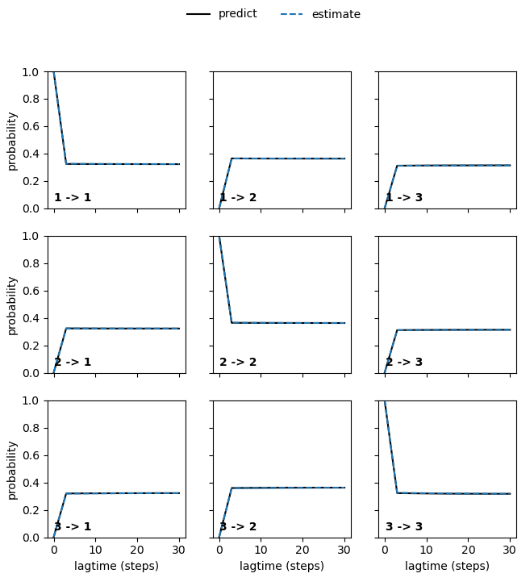


**(a)** Triple-well (connected) MSM Chapman-Kolmogorov test.

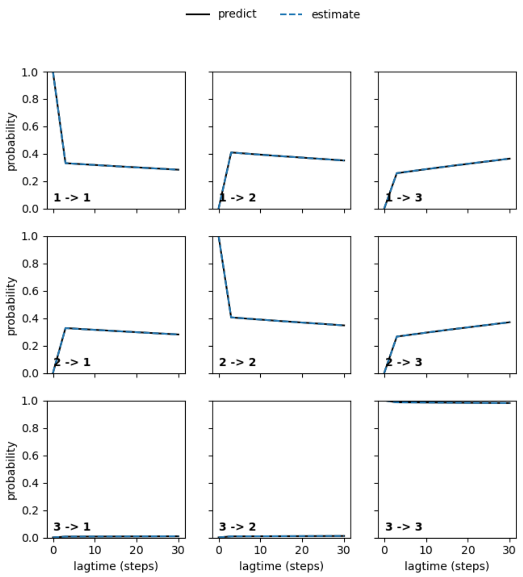


**(b)** Triple- well (missing) MSM Chapman-Kolmogorov test.

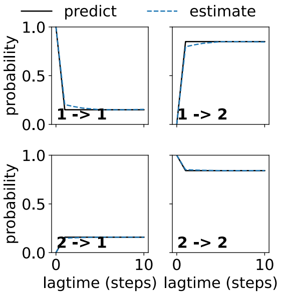


**(c)** Triple-well (disconnected, minor state) MSM Chapman-Kolmogorov test.

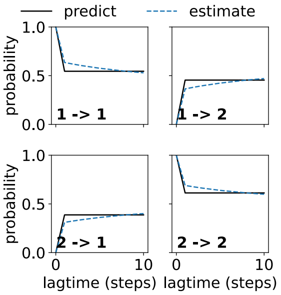


**(d)** Triple-well (disconnected, major states) MSM Chapman-Kolmogorov test.

**Fig. S2.** Triple well potential CK tests of all MSMs: connected (a), missing state (b) and disconnected (c and d).

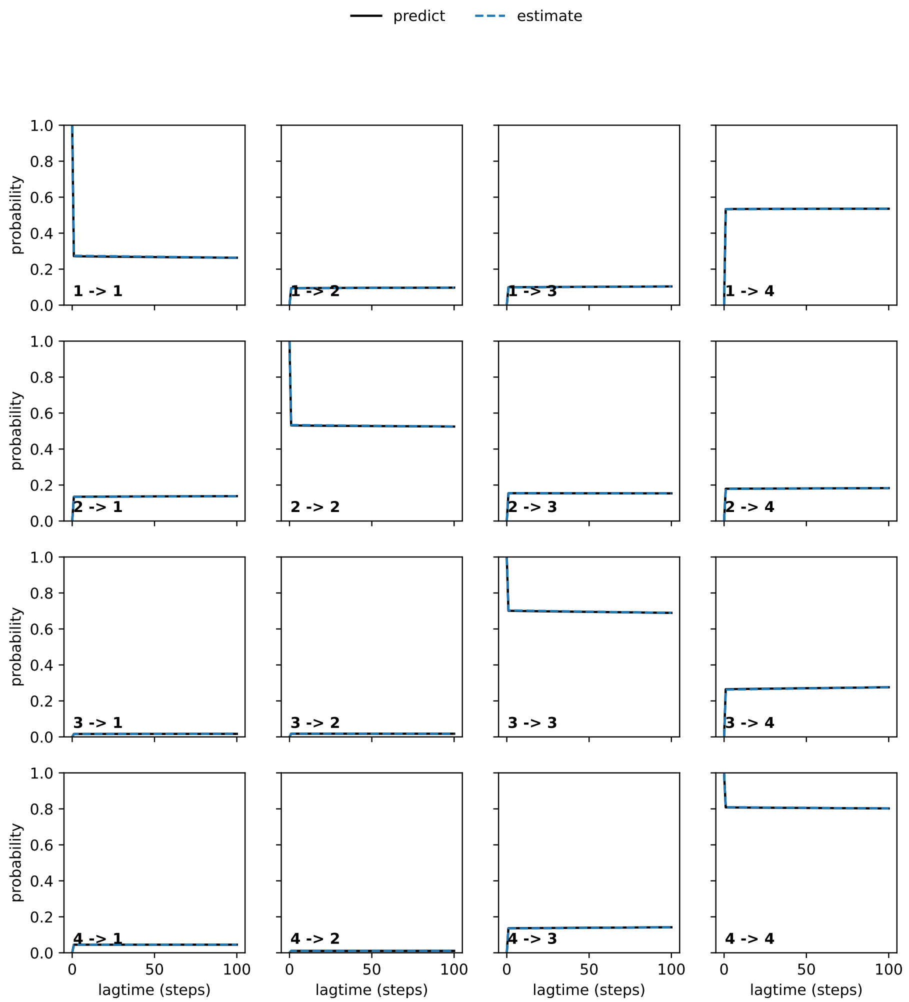


**(a)** BPTI (connected) MSM Chapman-Kolmogorov test.

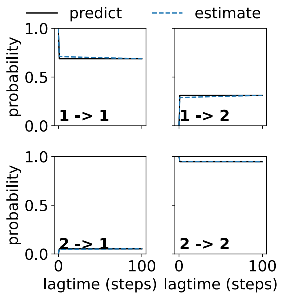


**(b)** BPTI (disconnected, major states) MSM Chapman-Kolmogorov test.

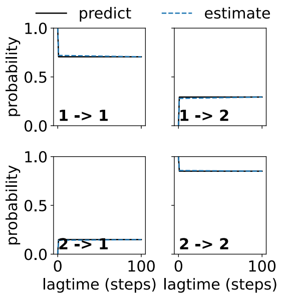


**(c)** BPTI (disconnected, minor states) MSM Chapman-Kolmogorov test.

**Fig. S3.** BPTI CK tests of all MSMs: connected case (a) disconnected case (b and c).

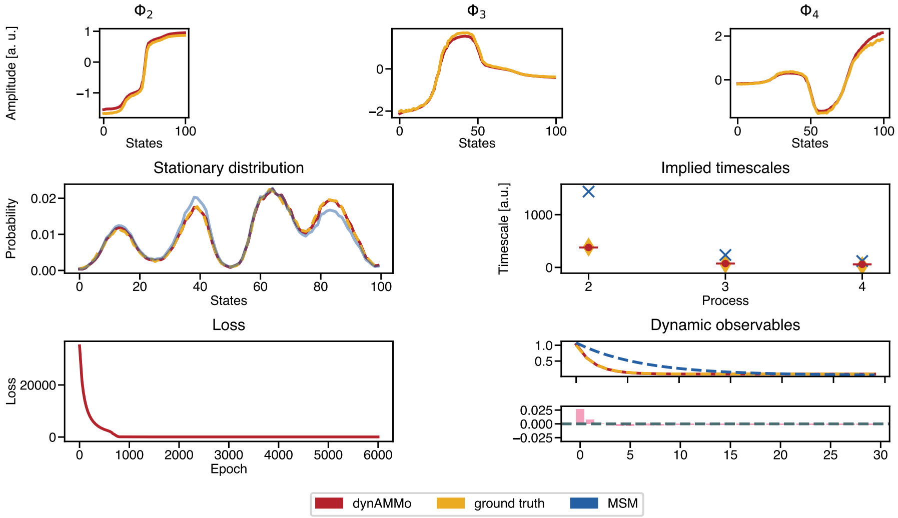


**(a)** Overview of Prinz potential connected scenario results.

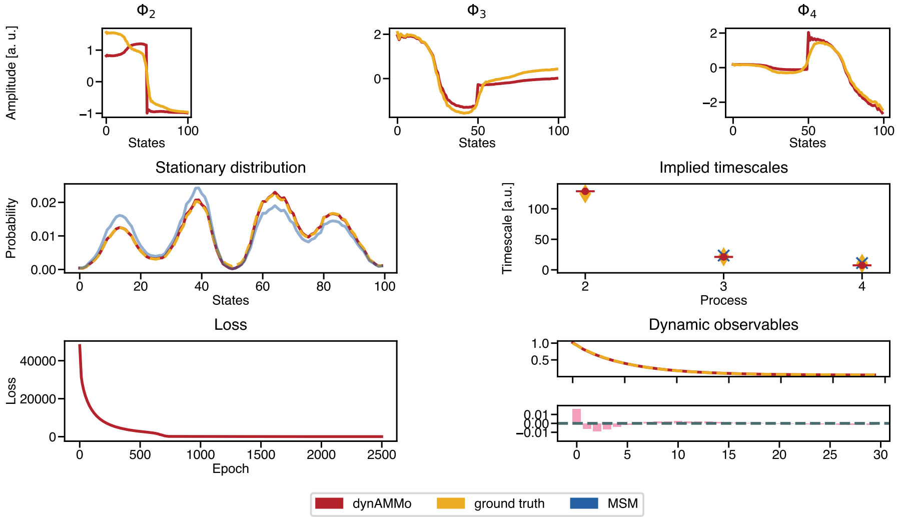


**(b)** Overview of Prinz potential disconnected scenario results.

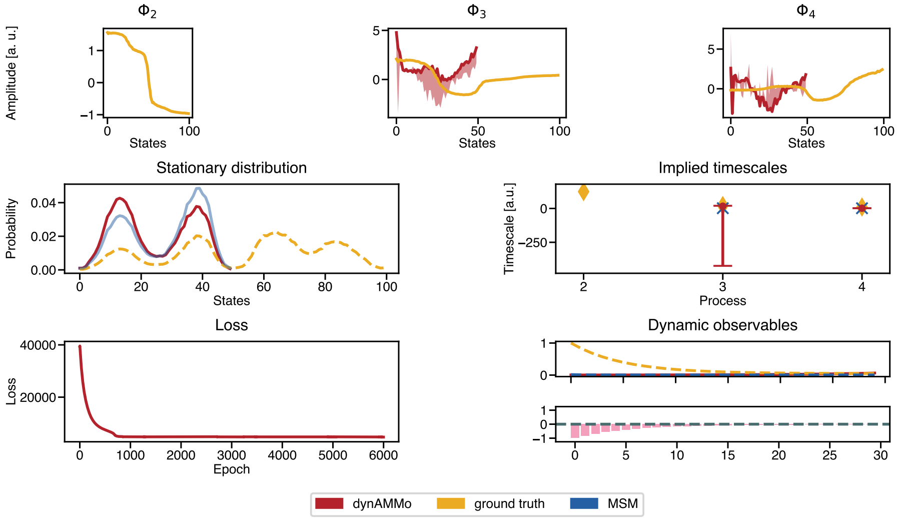


**(c)** Overview of Prinz potential unobserved-state scenario results.

**Fig. S4.** Comparison of Prinz potential datasets between dynAMMo (red), ground truth (yellow) and MSMs (blue). Top: the dominant eigenvectors are shown as a function of states. Middle left: stationary distribution as a function of states. Middle right: the timescales of the slowest processes are shown in the implied timescales plot. Bottom left: total loss as a function of epochs. Bottom right: The dynamic observable using the slowest eigenvector ($\mathbf{R}_2$) as observable function..

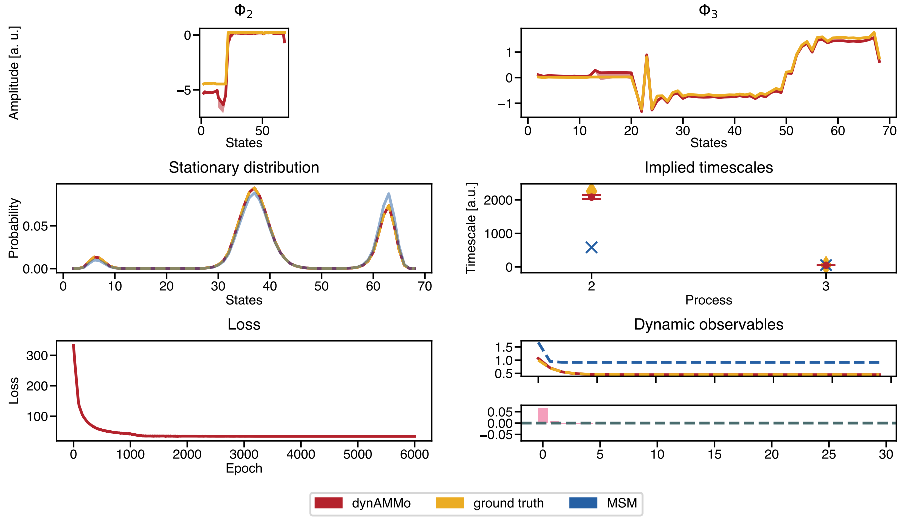


**(a)** Overview of triple well potential connected scenario results.

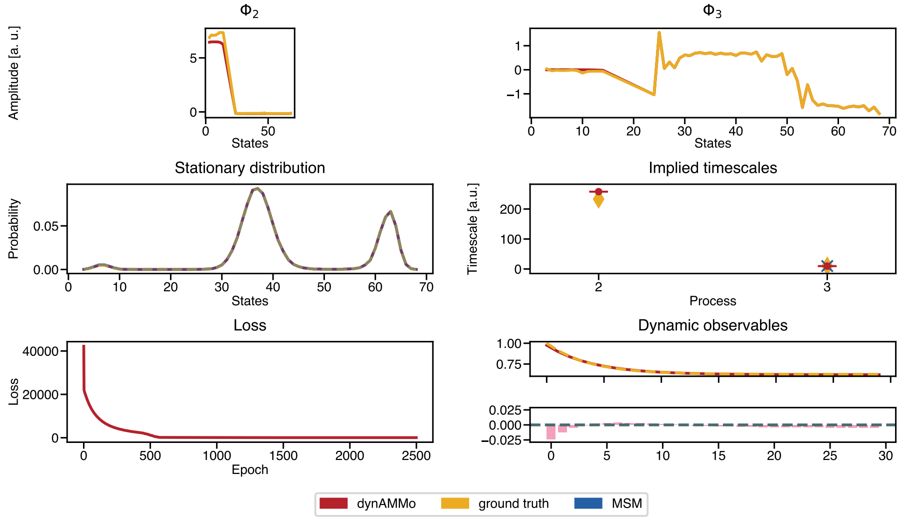


**(b)** Overview of triple well potential disconnected scenario results.

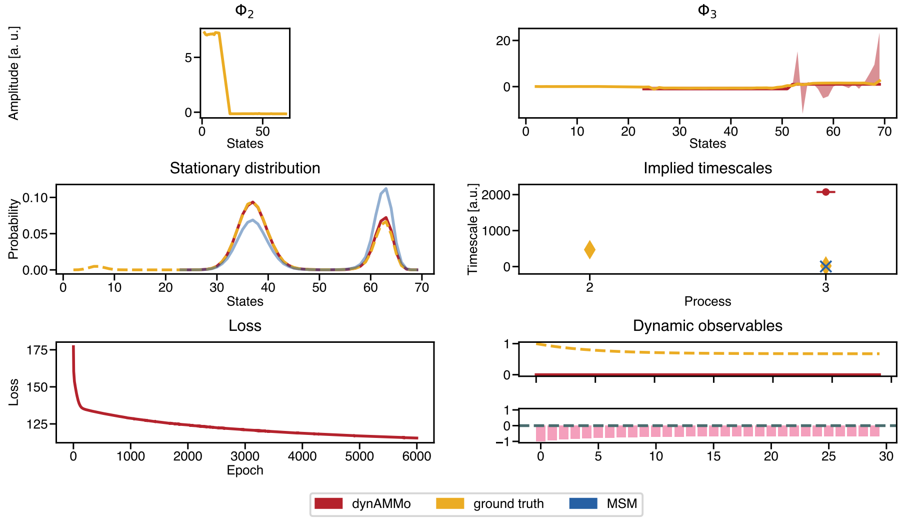


**(c)** Overview of triple well potential unobserved-state scenario results.

**Fig. S5.** Comparison of triple well potential datasets between dynAMMo (red), ground truth (yellow) and MSMs (blue). Top: the dominant eigenvectors are shown as a function of states. Middle left: stationary distribution as a function of states. Middle right: the timescales of the slowest processes are shown in the implied timescales plot. Bottom left: total loss as a function of epochs. Bottom right: The dynamic observable using the slowest eigenvector ($\mathbf{R}_2$) as observable function..

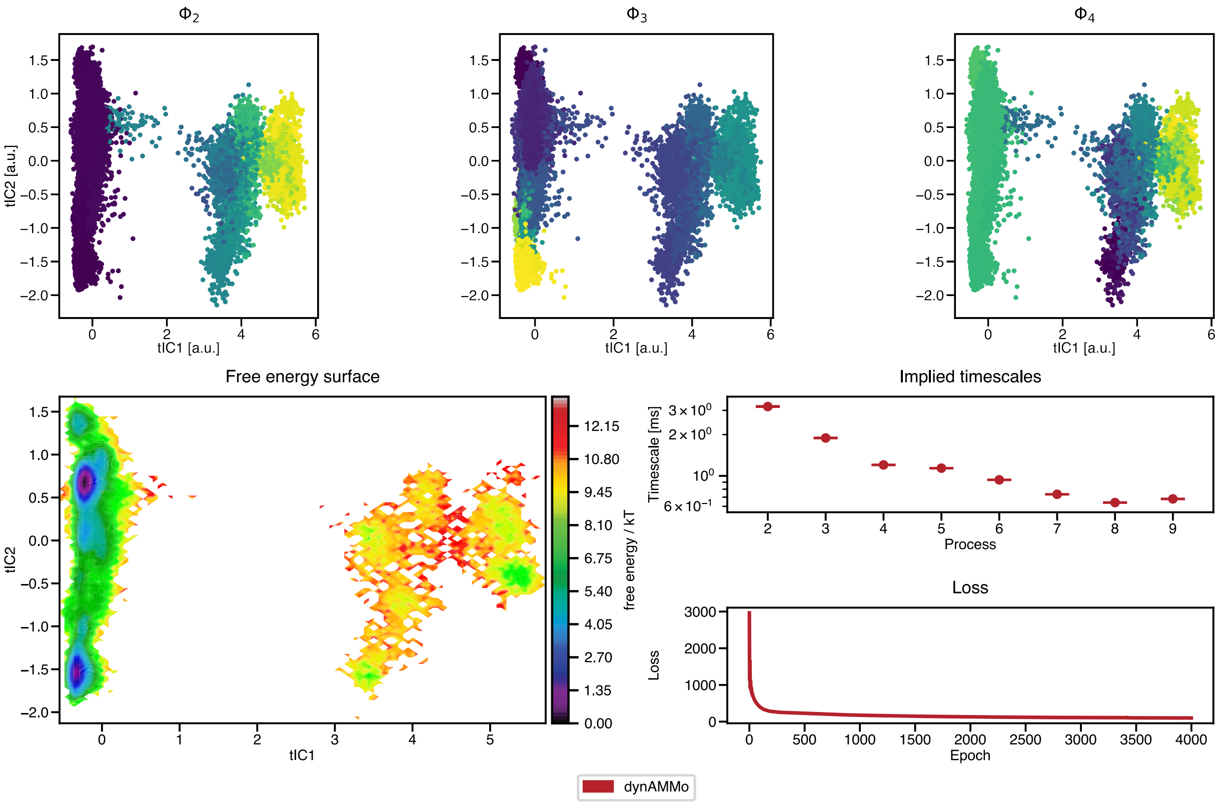


**(a)** Overview of BPTI connected scenario results.

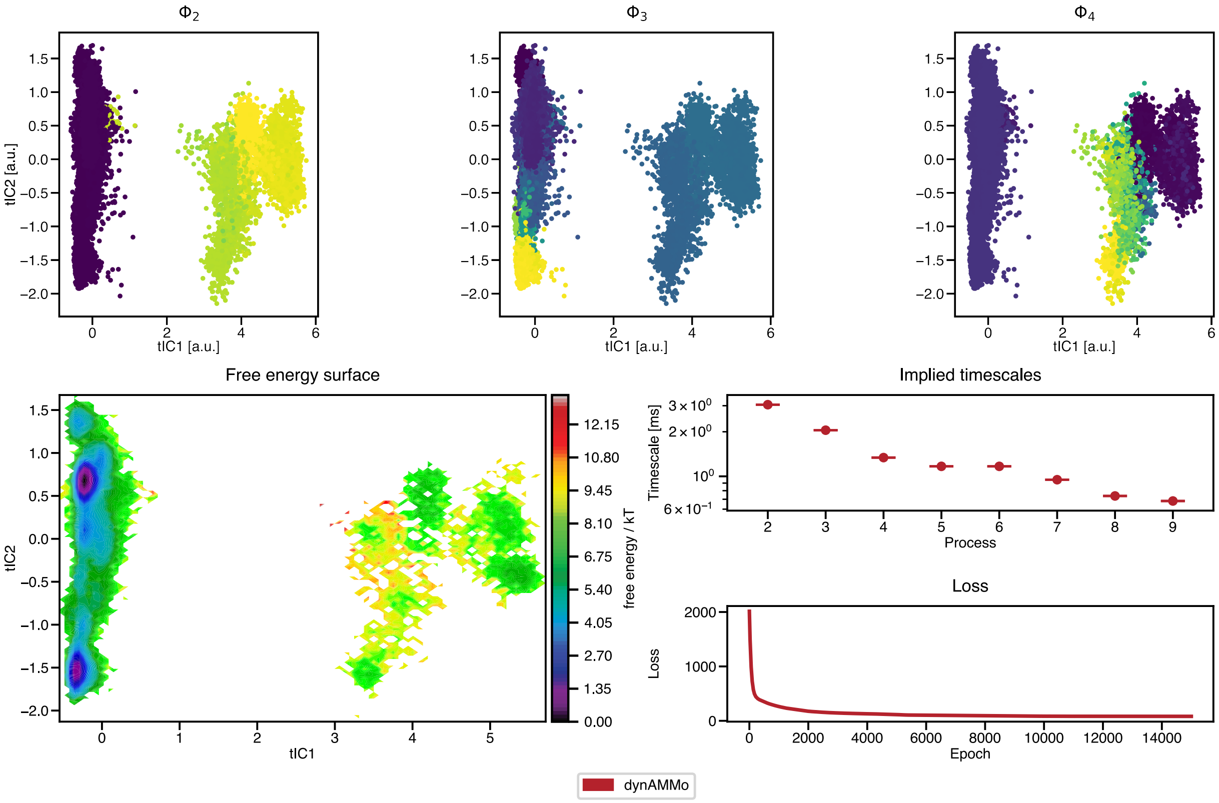


**(b)** Overview of BPTI disconnected scenario results.

**Fig. S6.** Comparison of BPTI datasets between dynAMMo (red), ground truth (yellow) and MSMs (blue). Top: the dominant eigenvectors are shown using the first and second time-lagged independent components. Bottom left: free energy surface of the first two tICs. Bottom right: implied timescales plot of the eight slowest timescales. Loss plot as a function of epochs.

| | Prinz potential | | | Triple-well potential | | |
|---|---|---|---|---|---|---|
| | biased | disconnected | missing | biased | disconnected | missing |
| dynAMMo | 0.55 ± 0.001 | 0.36 ± 0.003 | - | 2.96 ± 0.004 | 3.89 ± 0.003 | - |
| ground truth | 0.56 | 0.37 | 0.37 | 2.96 | 3.92 | 3.92 |
| MSM | 0.37 | - | - | 3.28 | - | - |

**Table S4. △G of slowest transition for the different model systems. All values are in $k_{\mathrm{B}}T$.**

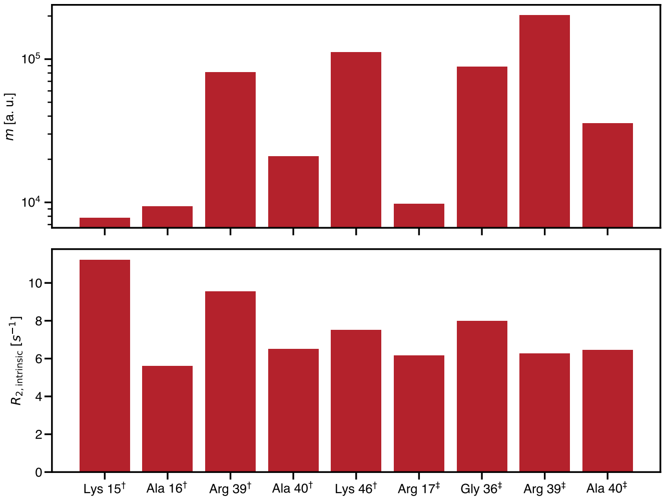


**Fig. S7.** Comparison of CPMG scaling parameters $m$ and $R_{2,\text{intrinsic}}$. The empirically determined scaling parameters for the eight residues shown in the main article.

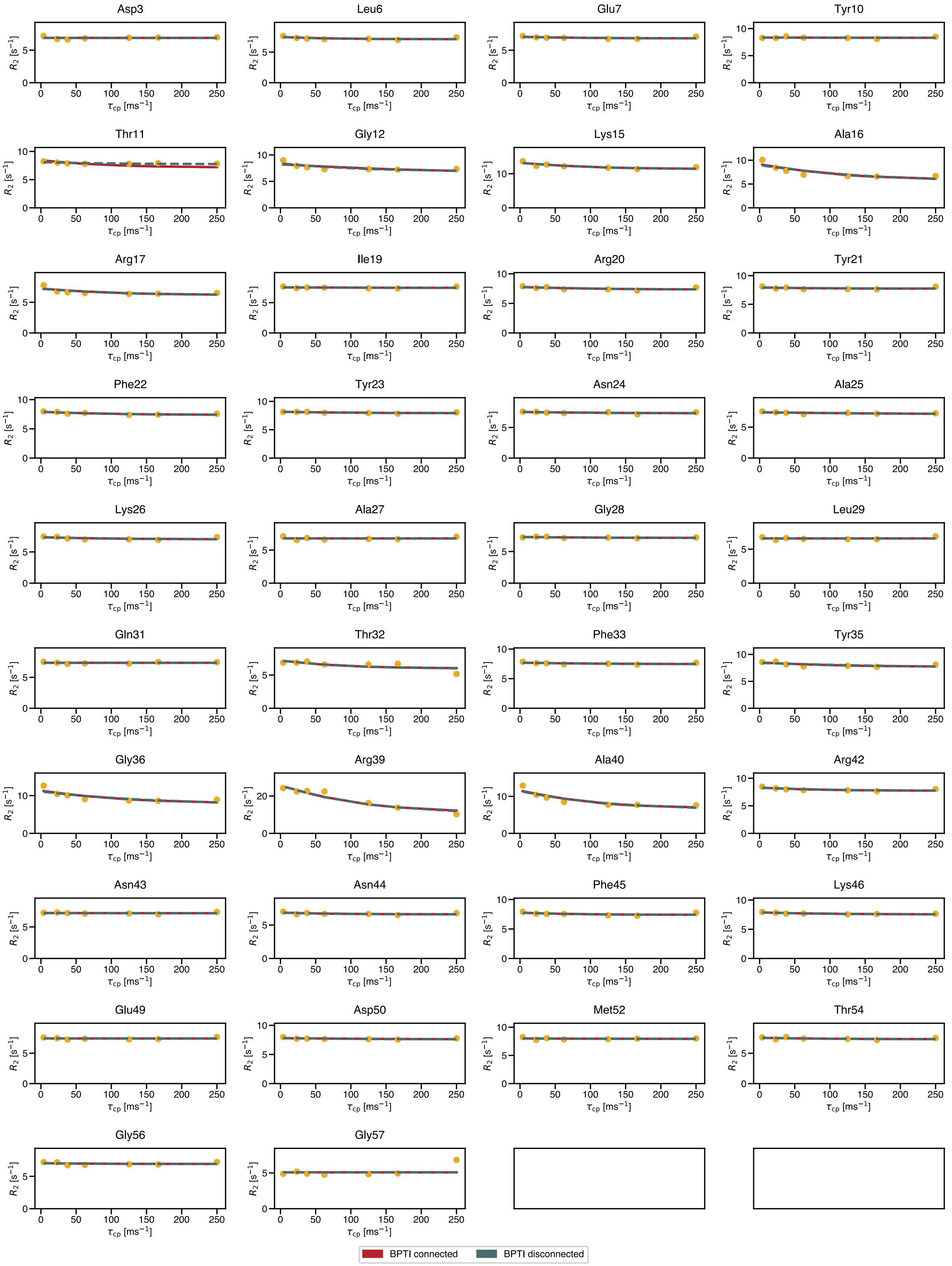


**Fig. S8.** Comparison of experimental BPTI CPMG data measured at 600 MHz with dynAMMo. The predictions of the two scenarios, connected (red) and disconnected (gray), are shown together with the experimentally acquired data (yellow, (14)).

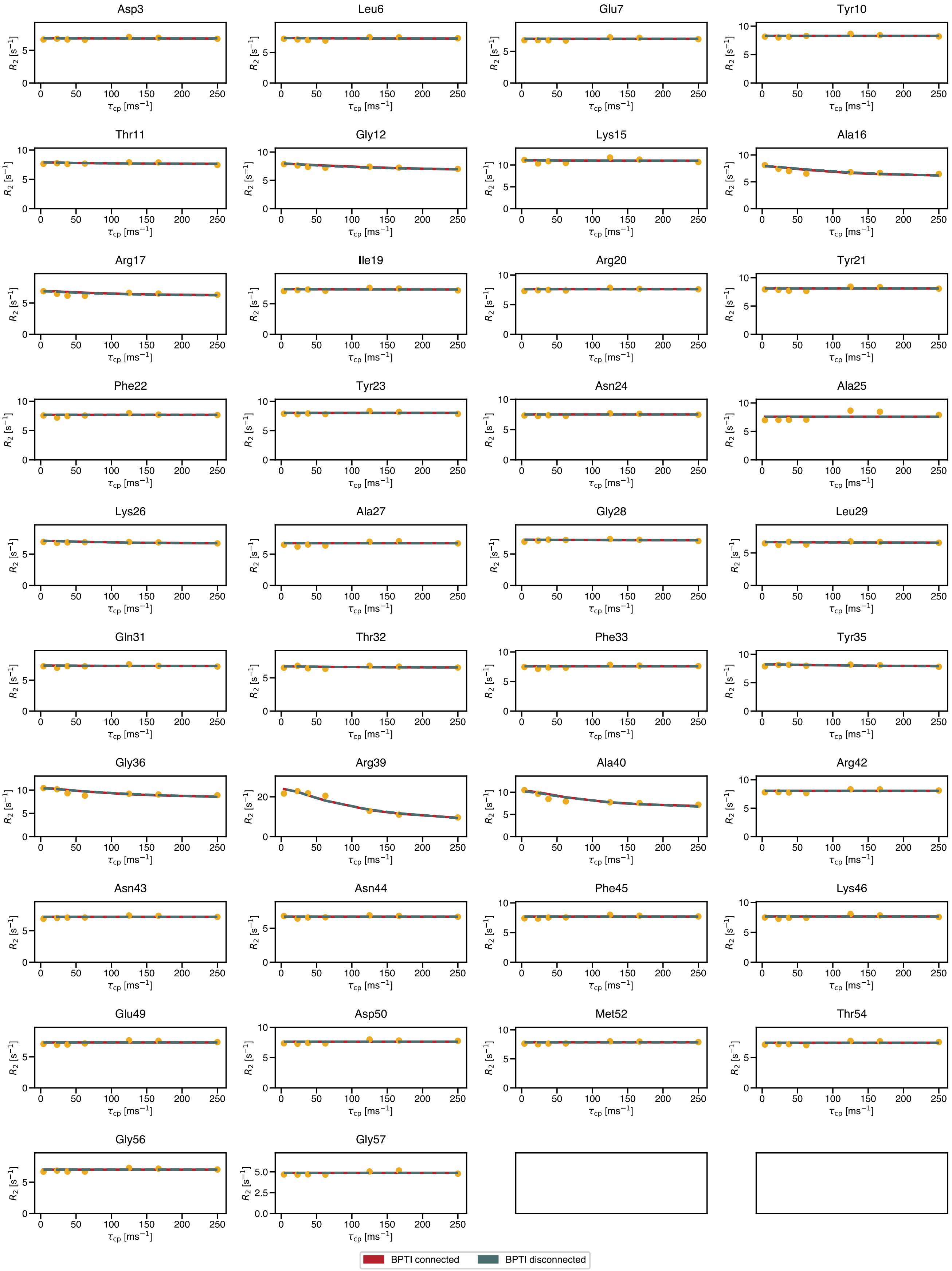


**Fig. S9.** Comparison of experimental BPTI CPMG data measured at 500 MHz with dynAMMo. The predictions of the two scenarios, connected (red) and disconnected (gray), are shown together with the experimentally acquired data (yellow, (14)).
